# VAMPIRE: Analyzing variation and motif pattern in tandem repeats

**DOI:** 10.1101/2025.06.15.659631

**Authors:** Zikun Yang, Shilong Zhang, Glennis A. Logsdon, Yafei Mao

## Abstract

Tandem repeats (TRs) are pervasive in eukaryotic genomes and play key roles in genome organization, evolution, and function, particularly in complex regions such as centromeres and subtelomeres. Although long-read sequencing technologies have improved the resolution of these regions, existing methods remain limited in their ability to systematically and accurately characterize large-scale TRs. Here, we introduce VAMPIRE, a k-mer–based computational tool for comprehensive TR discovery, annotation, and quantification. Unlike previous methods, VAMPIRE enables reference-free, fine-grained decomposition of both simple and complex TRs, capturing motif variation in sequence, length, and structure with high sensitivity and scalability. Applied to complete telomere-to-telomere (T2T) human and nonhuman primate (NHP) genome assemblies, VAMPIRE reveals previously unrecognized high-order repeat inversions within human centromeres—an underappreciated evolutionary mechanism contributing to centromere diversity. Additionally, the tool identifies lineage-specific and expanded TRs, including human-specific STR/VNTR expansions and NHP-specific subtelomeric heterochromatin (e.g., pCht/StSat), underscoring their dynamic turnover and structural complexity. VAMPIRE provides a robust and scalable framework for TR analysis in the era of long-read sequencing, with broad utility across human genetics, evolutionary biology, and the study of complex TRs in non-model organisms.

## Introduction

With the advance of long-read sequencing technologies, genome assemblies have achieved unprecedented levels of contiguity and accuracy^1^. It is now possible to generate complete, telomere-to-telomere (T2T) genome assemblies using multiple long-read sequencing technologies^2, 3^. These advancements have enabled the exploration of previously inaccessible genomic regions, including tandem repeats (TRs), segmental duplications (SDs), structurally divergent regions, and other complex genomic regions^4–7^.

TRs are widely distributed across eukaryotic genomes and exist in various forms, including short tandem repeats (STRs), variable number tandem repeats (VNTRs), and large satellite DNA arrays^8^, such as higher-order repeats (HOR) in centromeres (e.g., alpha-satellite) and subtelomeric heterochromatic caps (e.g., pCht/StSat sequences). These repetitive sequences are crucial in genome organization, regulation, evolution, and human diseases^9–11^. However, their characterization has long been hindered by the limitations of short-read sequencing and analytical methods, which failed to fully resolve their complexity^10, 12–14^. Despite the advent of nearly complete T2T genomes that now enable base-level resolution of TR regions, analytical frameworks capable of fully harnessing this resolution remain insufficiently developed.

To address the challenges of TR analysis, various tools have been developed, including TRF^15^, ULTRA^16^, uTR^17^, MotifScope^18^, VAMOS^19^, HumAS-HMMER_for_AnVIL^20^ and CENdetectHOR^21^ (Supplementary Table 1). While these tools have significantly advanced the field, most are designed for specific tasks and rely on additional tools or prior knowledge to complete the full process of TR analysis. For example, VAMOS is a reference-based tool based on StringDecomposer^22^ to determine human STR or VNTR haplotypes at the population level^19^, while HumAS-HMMER_for_AnVIL is an HMM-based method for annotating human centromeric satellites^20^. However, neither of these approaches is *de novo*; both require either a user-defined consensus motif or pre-trained HMM profiles. This reliance limits their effectiveness in comprehensive TR annotation and their applicability to non-model organisms. Additionally, many existing tools, like TRF, either lack the resolution needed to distinguish variations among motif types or struggle with analyzing large-scale TR arrays^15, 18^. Furthermore, these tools are generally incapable of detecting or interpreting complex structural patterns, such as inversions and transposable element (TE) insertions within TR arrays. These limitations can lead to misinterpretations of genomic structure and obscure critical biological insights.

Collectively, there remains a significant gap: no current solution can perform *de novo* motif discovery, decompose TR regions, and analyze associated mutations and structural variants (SVs). The development of such an integrated tool is essential for advancing our understanding of the biological significance, evolutionary dynamics, and functional roles of TRs in genomics.

Here, we introduce VAMPIRE, a tool designed to characterize **VAriation and Motif Patterns In tandem REpeats**, addressing the critical gap in TR analysis. VAMPIRE integrates *de Bruijn* graph construction, BK-tree indexing, edit distance refinement, and other algorithms to *de novo* identify motif types, characterize non-TR sequences, and decompose motif structures within TR regions. Benchmarking against both simulated and empirical datasets demonstrates that VAMPIRE achieves robust performance across simple and complex TRs. Applying VAMPIRE to recently published T2T human and nonhuman primate (NHP) genomes^2, 3^ revealed novel biological insights into lineage-specific and expanded TRs, including human-specific expanded STRs and VNTRs, and NHP subtelomeric repetitive sequences.

## Results

### Overview of VAMPIRE

VAMPIRE accepts a single FASTA file as input and supports multi-threaded processing to efficiently annotate TRs, ranging from STRs to megabase-long satellite sequences, at single-base resolution. VAMPIRE consists of three modules: motif finding, iterative motif searching, and motif alignment and chaining.

To perform *de novo* annotation of TRs without prior knowledge, we first segmented the genome sequence into windows of 5 kbp with a sliding step size of 1 kbp. We then constructed k-mers (default: k=5 bp) for each window and used these k-mers to build a *de Bruijn* graph. From the graph, we extracted simple loops, each representing a unique motif type, with the minimum edge weight serving as an estimate of its copy number (Method). Finally, we aggregated motifs across all windows, ranked them by their copy numbers, filtered out low-frequency motifs based on a user-defined threshold, and generated a raw motif set (Figure 1a).

**Figure 1.**
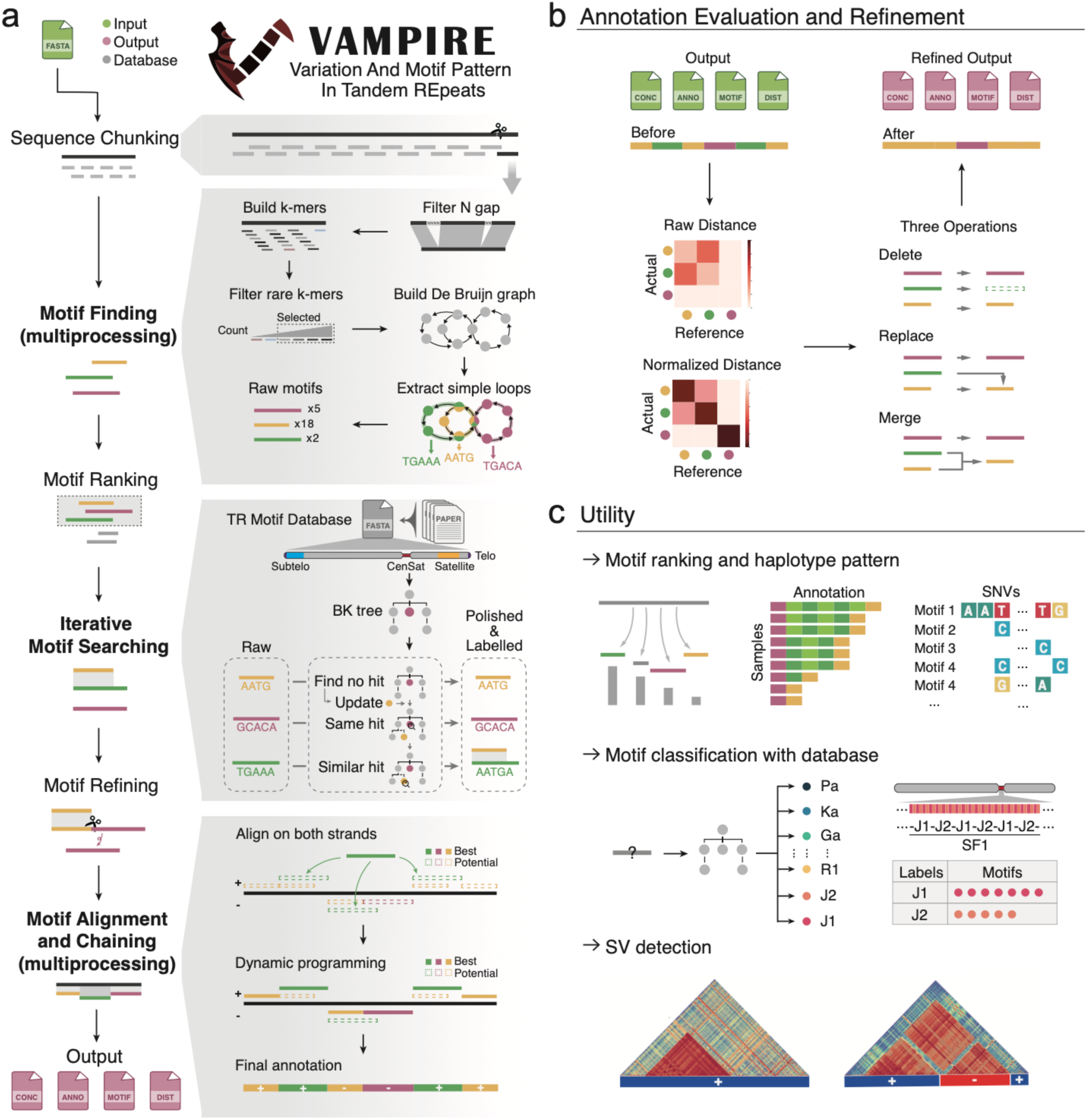
Overview of VAMPIRE algorithms and applications. (a) Schematic illustration of the three modules in VAMPIRE, including *de novo* motif discovery, iterative motif searching, and motif alignment and chaining. (b) Evaluation and global refinement of motif annotations. (c) Utility of VAMPIRE for motif annotation, classification, and structural variation characterization within TRs.

Due to the absence of a designated starting point within a loop, a single motif can appear in multiple equivalent forms (e.g., GGC, CGG, and GCG, which are identical, with the canonical form being GGC). These motif forms can obscure the underlying mutation patterns within the TR regions. To address this, we developed an iterative motif-searching module that identifies the canonical form for each motif, aligns similar motifs, and generates a high-quality, refined motif set (Figure 1a).

We utilized a BK-tree to efficiently identify the most similar match for a given motif type, enabling fast approximate searches based on edit distance. Optionally, a canonical motif database can be provided to initialize the BK-tree, after which raw motifs are progressively searched, starting from the most frequent and moving to the least frequent. If no similar match is found, the motif is treated as the canonical form and added to the tree for future comparison. When a match is found, two scenarios are considered: if the motifs are identical, no change is made; if they are similar but not identical, the existing canonical motif in the BK-tree serves as a reference to realign the new motif, which is then added to the tree (Figure 1a). This process ensures both the accuracy and efficiency of motif identification and standardization. If no canonical motif database is provided, the BK-tree is built directly from the raw motifs.

The optional database facilitates the alignment of *de novo* annotated motifs with established motif databases, ensuring consistency across studies. For instance, when annotating human centromeric HOR motifs, which already have standardized names, the database enables accurate mapping to these recognized nomenclatures, enhancing the reliability and coherence of our annotations.

To accurately identify complex motif structure in TR regions, we developed a motif alignment and chaining module. This module aligns the refined motif set to the genomic sequence and applies a dynamic programming algorithm to enable accurate quantification of motif copy numbers (Figure 1a). Meanwhile, unlike traditional approaches that assume motif contiguity, our module accommodates complex patterns, including non-TR sequence insertions and motif inversions.

Currently, our tool primarily focuses on annotating motifs and classifying the differences across them, but it lacks an evaluation of the annotation quality at the global level. To address this, we introduced a distance matrix approach to quantify the discrepancy between the actual sequence and the annotated motifs at the global level (Figure 1b, Supplementary Figure 1, Method). We implemented three operations—motif deletion, replacement, and merging—that enhance the accuracy of the motif annotation. These operations enable the removal of low-confidence motifs, compression of the motif set, and refinement of the global motif annotation.

In summary, our tool–VAMPIRE–can *de novo* identify motifs, rank them by copy numbers, and characterize variations (e.g., SNVs) across motifs using unified standardization. Additionally, when a known motif database is available, it can link the *de novo* motifs to established references. Moreover, since our annotation files include motif strand information and non-TR sequence insertion breakpoints, the tool is capable of identifying SVs (e.g., TE insertions and inversions) and motif patterns within the entire TR sequence region.

### Performance benchmarking of VAMPIRE and other TR detection tools on simulated data

To evaluate the performance of our tool and other TR detection tools—TRF^15^, uTR^17^, ULTRA^16^, and MotifScope^18^—in characterizing motif variations, non-TR insertions, and motif inversions, we generated simulated TR sequences with diverse motif types, lengths, and non-TR insertions using a custom TR generator (Methods, Supplementary Figure 2).

Non-TR sequences, such as TEs and SDs, are known to be inserted as spacers within TRs^3^. For instance, TEs are frequently observed in centromeric regions^4^, and SDs are present in subtelomeric heterochromatic regions of *Pan* and gorilla genomes^3, 23, 24^. In this study, we simulated 2,000 TR sequences with varying numbers of TE insertions (ranging from 1 to 3) and motifs containing 1% mutations (Figure 2a). Our results showed that VAMPIRE achieved the highest performance, detecting 81.3% of motifs and TE insertions on average (Figure 2b, c, Supplementary Table 2). MotifScope also demonstrated good performance, with a detection rate of 53.5%, but has a higher false positive rate to misannotate TE sequences as TRs (Supplementary Figure 3). Although TRF was able to identify all TRs, it failed to detect TE insertions in 83.1% of simulated sequences due to its reliance on consensus sequences rather than detecting insertions. ULTRA and uTR, on the other hand, were unable to detect TRs when motifs contained a certain mutation rate.

**Figure 2.**
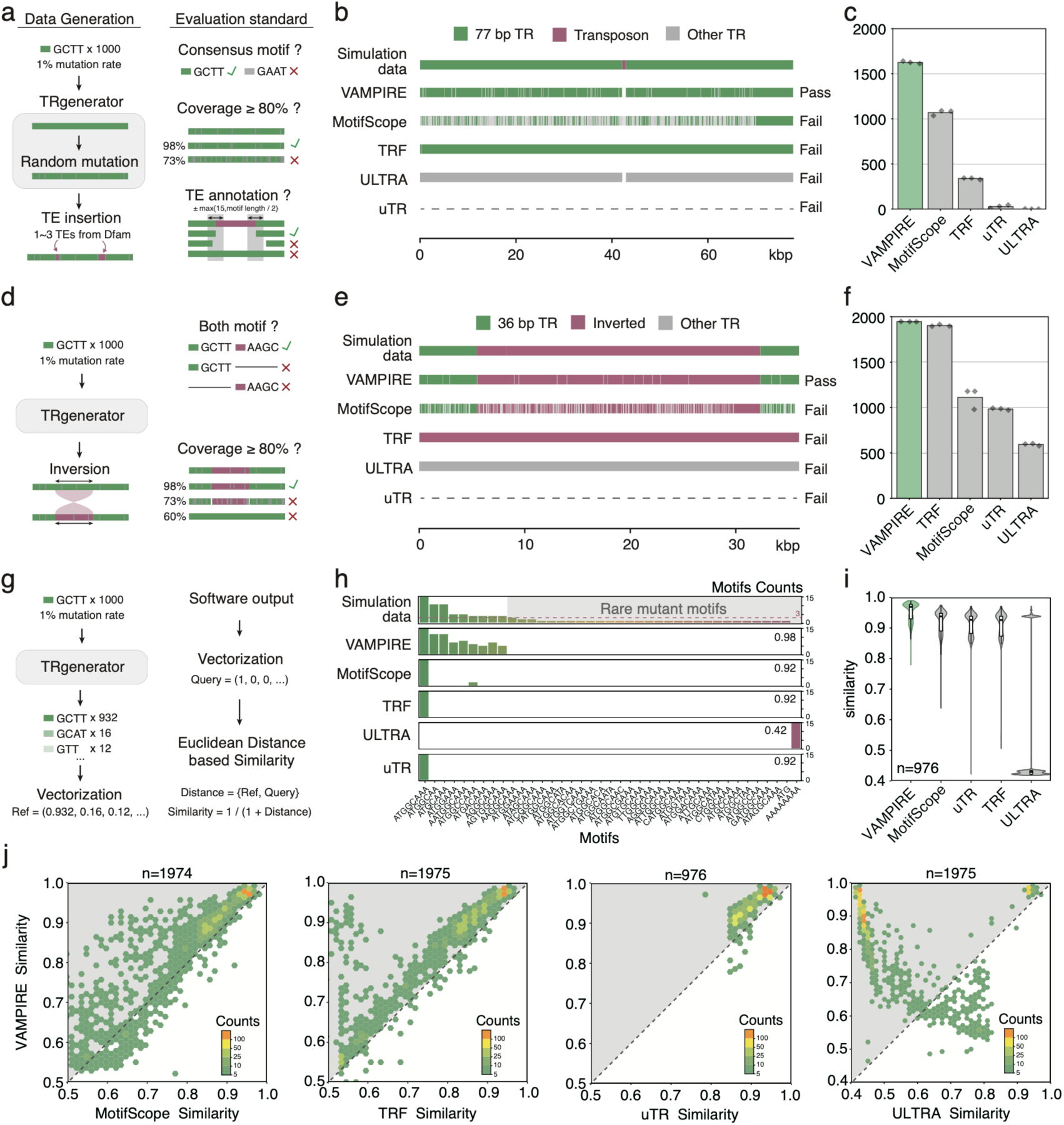
Benchmarking TR annotation using simulated datasets. (a–c), Evaluation of transposable element (TE) insertions within tandem repeat (TR) sequences. (a) Schematic overview of the simulation strategy and evaluation criteria for TE insertions within TRs. (b) Representative examples of simulated TR sequences with TE insertions classified as pass or fail. (c) Number of correctly detected TE insertions by each tool across three independent simulation replicates. (d–f) Evaluation of inverted motif detection. (d) Schematic of TR sequences containing both original and reverse complement motifs, along with the evaluation standard. (e) Representative examples of successful and failed inversion motif detection. (f) Number of correctly identified inversions by each tool across three simulation replicates. (g–i) Assessment of motif variation detection. (g) Illustration of simulation strategy for TR sequences with variable motif types and evaluation standard. (h) Distribution of detected motif types across tools. (i) Motif similarity distribution between annotated motifs and ground-truth motifs for each tool, highlighting detection accuracy. (j) Scatter plot comparing motif similarity scores between the simulated reference and tool annotations. The Y-axis shows similarity between VAMPIRE and the simulated distribution; the X-axis shows similarity for other tools. VAMPIRE shows higher concordance with the ground truth.

Motif inversion is another key factor contributing to the complexity of TRs^4^, particularly in satellite DNA arrays. For instance, a 1.7 Mbp inversion has been identified within the centromere of chromosome 1 in T2T-CHM13^4^. While most current tools either overlook inversions or annotate inverted regions as distinct motifs without considering motif strand orientation, we evaluated inversion detection by generating a benchmark dataset with inverted TR segments (Figure 2d, Method). Our results showed that both TRF and VAMPIRE outperformed other tools, with detection rates of 97.3% and 95.1%, respectively (Figure 2e, f, Supplementary Figure 3, Supplementary Table 2). In contrast, the remaining tools only detected inversions in 29.7% - 55.6% of the simulated sequences.

TR regions exhibit mutation rates significantly higher than simple genomic regions^25^, leading to diverse motif types within a TR region. However, many tools tend to overlook these mutations. To evaluate each tool’s ability to capture motif variants, we used a Euclidean-distance-based similarity metric to compare the motif frequency distribution between simulated data and tool-generated annotations for 2000 simulated TR sequences (Figure 2g). VAMPIRE can not only correctly detect the consensus motif but also capture the real distribution of mutation (Figure 2h, Supplementary Figure 3). Among the 976 sequences annotated by all five tools, VAMPIRE showed the best performance, with Euclidean similarity ranging from 78.0% to 98.9% (Figure 2i, Supplementary Table 3). Pairwise comparisons between VAMPIRE and the other four tools consistently showed that VAMPIRE annotated motif types/variations more accurately (Figure 2j). Specifically, VAMPIRE exhibited up to 3.51%, 4.16%, 5.19%, and 31.7% higher similarity on simulated data compared to MotifScope, uTR, TRF, and ULTRA, respectively.

### Empirical data application and novel insights

To validate the reliability and utility of VAMPIRE, we applied it to the T2T-CHM13 genome, with a particular focus on STRs, VNTRs, and alpha-satellites at centromeres. We aimed to assess whether VAMPIRE could recapitulate previously established TR motif annotations and to explore whether it could provide new insights beyond existing methods^15^.

Given that TRF has been a foundational tool for STR and VNTR consensus motif annotation^15^, we benchmarked VAMPIRE against TRF for these TRs. To ensure a fair and consistent comparison, we applied a unified set of filtering strategies (e.g., motif copy number >=5 and excluding centromere satellites, Method) to the raw TR annotations generated by both tools. Under these criteria, ∼94% of TR loci were consistently identified by both VAMPIRE and TRF (Figure 3a). Of the remaining ∼6% of loci not jointly detected, the majority contain large motif variations that exceed the threshold set in VAMPIRE (Supplementary Figure 4a-b). The discrepancy primarily arises from differences in sensitivity between the tools when determining motif length, variation, and copy number. For example, some TRs identified by VAMPIRE did not meet the joint filtering thresholds due to slight underestimation of repeat length or copy number by TRF (Supplementary Figure 4c). These results show that VAMPIRE performs robustly in annotating STRs and VNTRs in the human genome.

**Figure 3.**
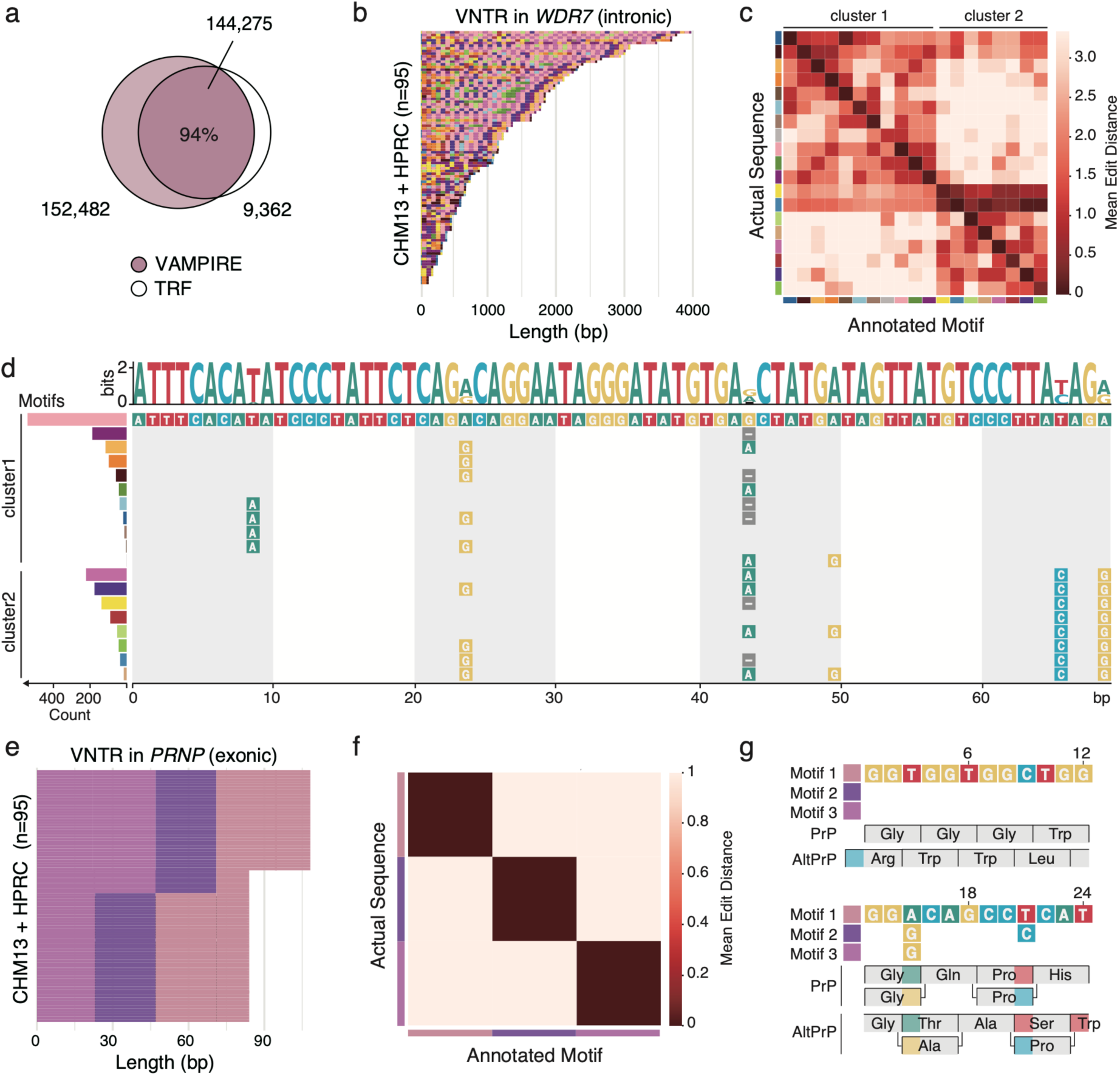
Characterization of simple TRs in human genomes using VAMPIRE. (a) Venn diagram showing 94% concordance in simple TR annotations (STRs and VNTRs) between VAMPIRE and TRF on the T2T-CHM13 assembly. (b) An example VNTR within the intronic region of WDR7 (chr18:57,226,379–57,227,527, T2T-CHM13), showing variable copy numbers and multiple motif types, each represented by a distinct color. (c) Heatmap of edit distances between annotated motifs and the corresponding genomic sequences shows high annotation accuracy by VAMPIRE and separation into two motif clusters. (d) Sequence logos of the two motif clusters in panel c highlight distinct sequence variation. (e) A VNTR within the exonic region of PRNP (chr20:4,738,616–4,738,701, T2T-CHM13) shows variable copy numbers of three different motifs across 95 long-read human genomes. (f) Edit distance heatmap for the PRNP VNTR motif shows clear separation between the three motif types. (g) The three motif types potentially alter the alternative reading frame (AltPrP) of PRNP. While the canonical PrP sequence remains unaffected, motif 2 and motif 3 may change the AltPrP peptide from GLY–THR–ALA–SER to GLY–ALA–ALA–PRO.

Beyond consensus motif detection, VAMPIRE excels in annotating motif variation and copy number variation in the human population. To illustrate this, we highlighted two examples of VNTRs within *WDR7* and *PRNP*. The TR located in an intron of *WDR7* (chr18:57226379– 57227527 in T2T-CHM13) consists of a 69 bp motif exhibiting extensive sequence variation and highly variable copy numbers across the human population (Figure 3b, Supplementary Figure 5). Using VAMPIRE, we *de novo* identified 19 distinct motifs, which could be broadly grouped into two clusters based on distance matrix analysis (Figure 3c, Supplementary Table 4). Within the 69 bp consensus motif sequence, four positions (24th, 44th, 66th, 69th) showed substantial variability, with the two clusters differing primarily at two of these sites (66th, 69th) (Figure 3d, Supplementary Table 5). These findings are consistent with previous studies^26^ and provide enhanced resolution of motif variation and pattern.

Another VNTR case involves the expansion of a 24 bp motif within the exon of *PRNP* (chr20:4738616–4738701 on T2T-CHM13). *PRNP* encodes two isoforms: the canonical PrP protein, which plays a critical role in neuroprotection, cell signaling, and synaptic maintenance, and an alternative isoform, AltPrP, which has distinct structural and potentially pathogenic properties^27, 28^. Overexpansion of the VNTR region within *PRNP* has been associated with an increased risk of prion diseases, including Creutzfeldt-Jakob disease and other transmissible spongiform encephalopathies^29, 30^.

Using VAMPIRE, we achieved a zero-base-error annotation of *PRNP* VNTR and identified three distinct motif variants, differing at the 15th base (A/G) and the 21st base (A/T) (Figure 3e, f, Supplementary Figure 6, Supplementary Table 6). Interestingly, these nucleotide substitutions do not alter the amino acid sequence of the canonical PrP isoform; however, they do result in amino acid changes in the alternative AltPrP isoform (Figure 3g). These findings extend our understanding of how motif-level variations within VNTRs can contribute to protein diversity and potentially disease risk at the population level.

In addition to annotating small TRs, we applied VAMPIRE to systematically characterize the active HOR regions of all chromosomes (22+XY) in the T2T-CHM13 genome^2^ (Supplementary Table 7). The sequence similarity matrices generated by VAMPIRE closely mirrored those from ModDotPlot^31^ across all chromosomes without considering motif inversions (SSIM n=26, Mean=0.97, s.d.=0.012) (Figure 4a-b, Supplementary Figure 7-9), validating the robustness of our approach. Beyond these, VAMPIRE additionally detected 6 chromosomes containing 17 TE insertions within the active HOR regions (Supplementary Figure 9, Supplementary Table 7).

**Figure 4.**
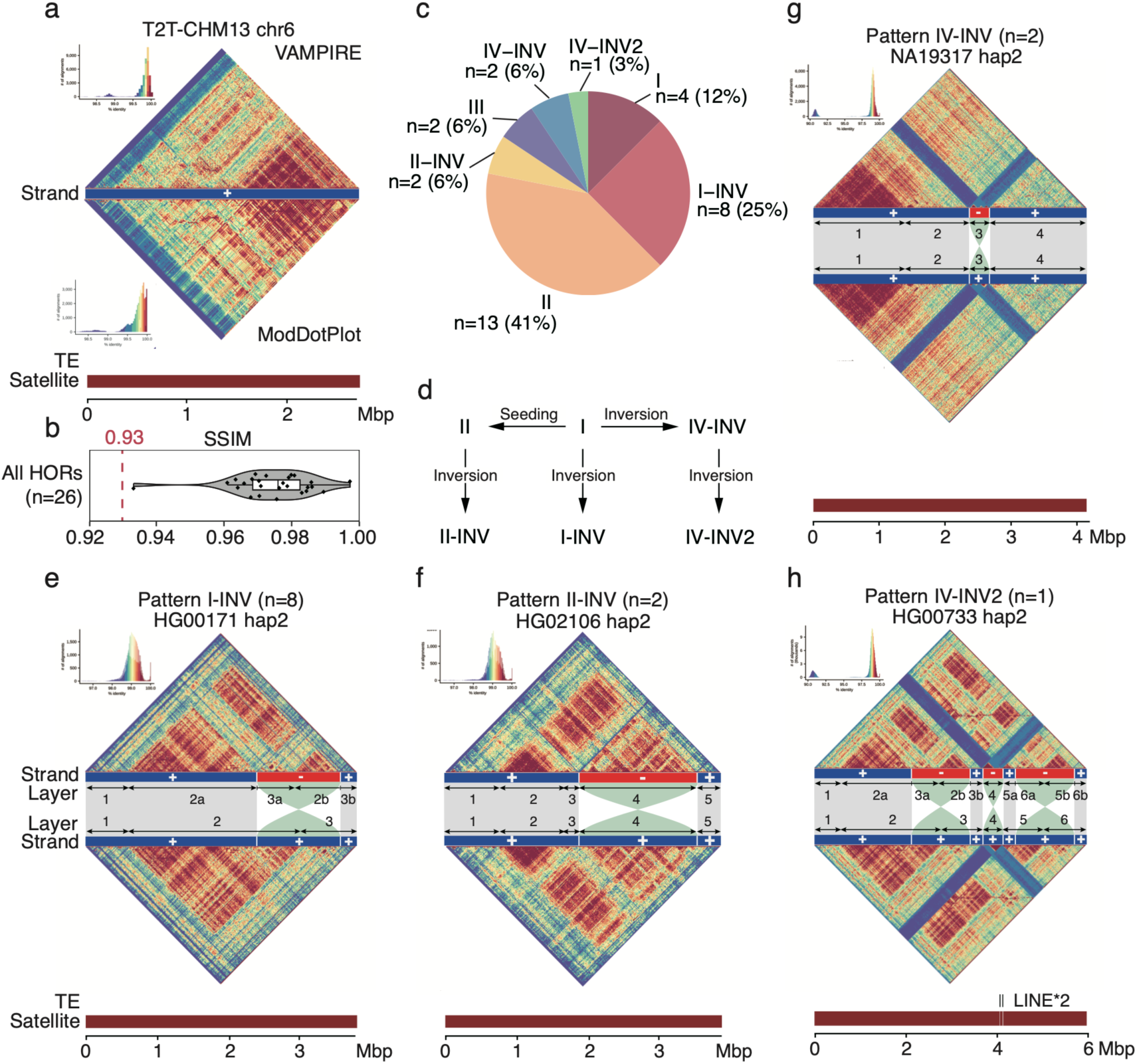
Centromere structure and inversion analysis using VAMPIRE. (a) Comparison of the active HOR region on human chromosome 6 reveals high concordance between results generated by VAMPIRE and ModDotPlot. Dark red shading in the satellite track denotes alpha-satellite sequences. (b) Structural similarity index (SSI) statistics show that VAMPIRE’s similarity heatmaps closely replicate those from ModDotPlot across all 26 active HOR regions analyzed. (c) Pie chart illustrating the distribution of seven distinct centromeric structural patterns observed on chromosome 1 across 32 human genomes. Three of these patterns (n = 13) include motif inversions. (d) Schematic model shows the proposed evolutionary trajectory of centromeric structural pattern diversification. (e) Similarity heatmap of Pattern I-INV. Top: observed matrix showing an inversion event. Bottom: artificially corrected matrix after inverting the inverted motifs. The inversion splits a previously homogenized motif block into two distinct regions, suggesting that unaccounted inversions can mask underlying HOR structural homology. (f) Similarity heatmap for Pattern II-INV. The inversion maintains motif distinctiveness from surrounding HORs, likely due to reduced motif homogenization caused by inversion. (g) Similarity heatmap for Pattern IV-INV, showing a comparable effect of inversion-driven motif isolation. (h) Another similarity heatmap for Pattern IV-INV, showing that an older central inversion (as in g) is flanked by two additional, more recent inversion events, which further disrupt previously homogenized motif blocks and reveal centromere diversity.

We next focused on human chromosome 1 and applied VAMPIRE to 32 T2T human assemblies in the HGSVC3 dataset^32^. Again, VAMPIRE’s results are highly similar to those from ModDotPlot^31^ (SSIM n=32, Mean=0.97, s.d.=0.005). Across these assemblies, VAMPIRE identified 15 distinct inversion events in 13 individuals (Supplementary Tables 8 and 9). We defined seven centromeric structural patterns based on the presence or absence of inversion events and the organization of sequence similarity blocks (Figure 4c-f, Supplementary Figure 10-14). Pattern I (n = 4) represents the most structurally simple configuration, characterized by the absence of inversions and the presence of a single, contiguous block of high sequence similarity (Supplementary Figure 13). Pattern II (n = 13) similarly lacks inversions but exhibits multiple high-similarity blocks interspersed with older, more diverged segments (Supplementary Figure 13). In contrast, Pattern III (n = 2) comprises multiple high similarity blocks with distinct sequence composition, suggesting independent expansion events of alpha-satellites (Supplementary Figure 13). These three patterns likely reflect varying degrees of TRs and sequence homogenization in the absence of inversion.

Inversions introduce additional complexity into centromeric architecture. Pattern I-INV (n = 8) likely derives from Pattern I through an inversion encompassing both the active HOR array and adjacent inactive regions, resulting in fragmentation of the previously contiguous similarity block (Figure 4e, Supplementary Figure 14). Pattern II-INV (n = 2) contains inversions restricted to the active HOR array and may have evolved from Pattern II via more localized rearrangements (Figure 4f, Supplementary Figure 14). Pattern IV-INV (n = 2) features inversions confined to the inactive flanking regions, while Pattern IV-INV2 (n = 1) appears to represent a derivative state, marked by two additional, more recent inversion events (Figure 4g, h, Supplementary Figure 14).

In the absence of inversion-aware analysis, conventional sequence similarity heatmaps from Moddotplot^31^ or StainGlass^33^ are unable to distinguish among the Pattern I-INV, II, and II-INV configurations (Supplementary Figure 12), potentially leading to erroneous interpretations of centromeric evolutionary history. Although these three patterns exhibit superficially similar architectures—each consisting of three inactive regions flanked by two active HOR arrays—the underlying evolutionary origins and mechanisms are distinct.

In Patterns II and II-INV, the expansion of Block 2 and Block 4 is likely driven by a motif “seeding” mechanism^34^. In contrast, what appear to be distinct blocks—Block 2a and Block 2b—in Pattern I-INV are derived from a single ancestral Block 2 prior to the inversion. Similarly, the separated Block 3a and Block 3b originate from the same ancestral HOR Block (Block 3), suggesting that their apparent divergence is an artifact of the inversion event acting on older, homologized repeat units. This reinterpretation underscores the importance of incorporating inversion-aware frameworks to accurately resolve structural relationships within centromere HOR structures.

Moreover, in Pattern IV (n = 2, e.g., NA19317_hap2), we identified a large inversion within the inactive HOR array that exhibited markedly reduced sequence similarity relative to other HOR regions—likely reflecting asymmetric motif homogenization between forward and reverse strands. A similar inversion was also observed in Pattern IV-INV (n = 1, HG00733_hap2), which harbors the lowest overall sequence similarity across all HORs analyzed. This inversion is further flanked by two additional, younger inversion events, suggesting a complex evolutionary history shaped by multiple structural rearrangements.

Together, these findings illustrate a continuum of inversion-mediated architectural variation in human centromeres. VAMPIRE’s ability to resolve these fine-scale features enables the dissection of structurally similar but mechanistically distinct centromere configurations, providing new insights into the evolutionary dynamics and diversity of centromeric regions in the human genome.

### New biology from lineage-specific TR expansion in primate evolution

Previous studies on human-specific TR expansions primarily relied on “squashed” long-read assemblies and the GRCh38 reference genome^10^, which likely resulted in incomplete or biased TR expansion catalogs. Here, we applied VAMPIRE to newly sequenced, high-quality phased human genomes^32^ (n=20) and T2T NHP genomes^3^ (n=11) to systematically re-examine human-specific TR expansions. Out of 618,719 TR loci on autosomal chromosomes and chromosome X, we identified 0.5% (n=3,100) human-specific TR expansions (HSEs) (Supplementary Table 10). Among these, 53.8% and 44.9% are located in intergenic and intronic regions, respectively (Figure 5a-d). To investigate how HSEs emerged during ape evolution, we analyzed their copy number profiles across nonhuman primates. We found that 68.5%, 60.0%, 75.0%, and 68.0% of the HSE loci located in intergenic, exonic, UTR, and intronic regions, respectively (Figure 5e-h, Supplementary Figure 15a), exhibit relatively stable copy numbers in nonhuman primates but show a sharp increase specifically in the human lineage (hereafter referred to as “burst“; see Methods). This pattern suggests that many HSEs likely originated from lineage-specific bursts at ancestral stable TR loci, rather than from a gradual expansion process that simply accelerated in humans.

**Figure 5.**
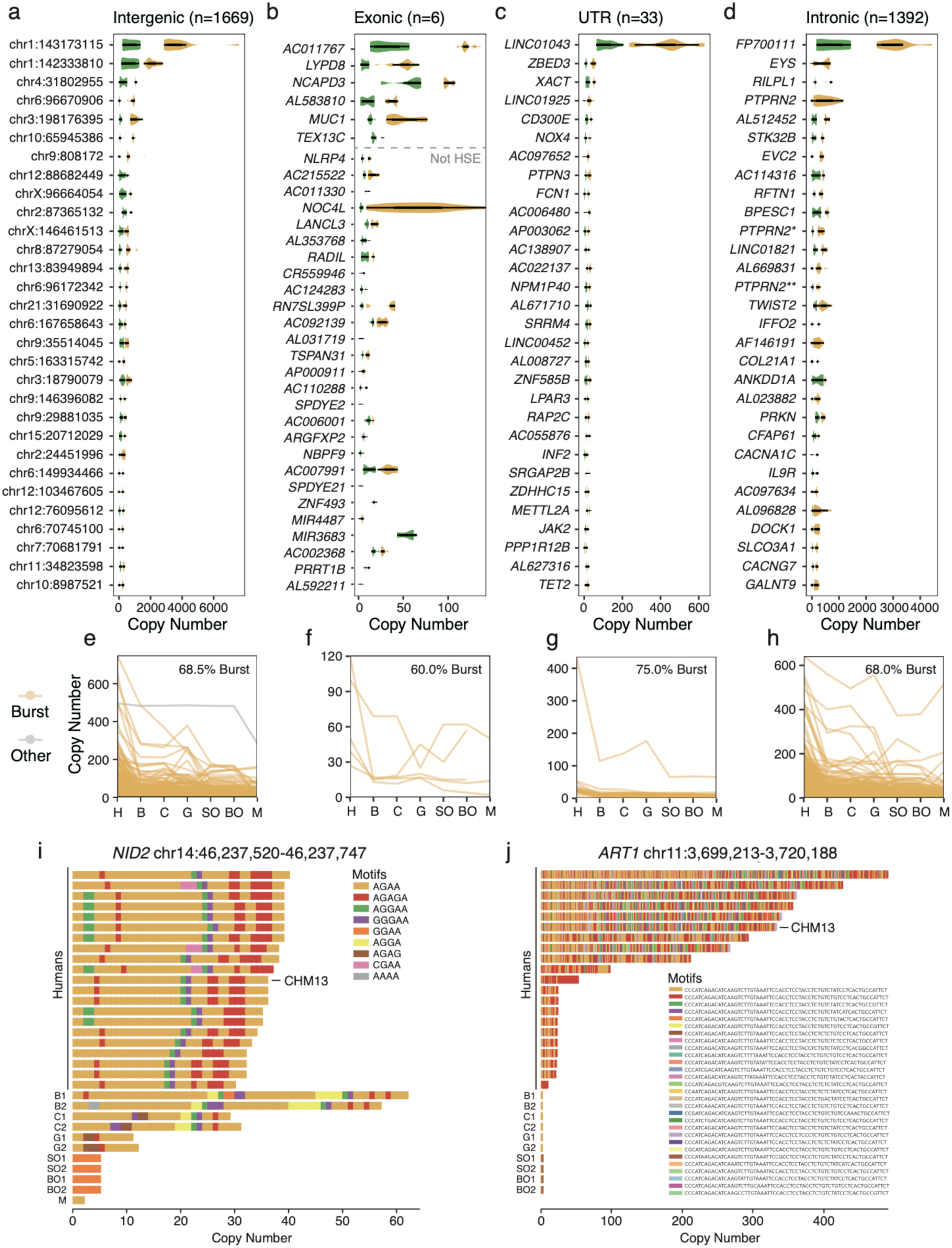
Human-specific expanded tandem repeats (HSE TRs) and their copy number dynamics. The Top 30 HSE TRs located in intergenic (a), exonic (b), UTR (c), and intronic (d) regions. Average copy numbers across the human lineage and nonhuman primates are displayed for intergenic (e), exonic (f), UTR (g), and intronic (h) regions, with loci exhibiting burst expansion in humans highlighted in yellow (Methods). Burst expansion in the human lineage is observed in 68.5%, 60.0%, 75.0%, and 68.0% of HSE TRs in the intergenic, exonic, UTR, and intronic regions, respectively. (i) With the availability of complete nonhuman primate assemblies, the previously reported intronic HSE STR locus in the *NID2* gene (identified in Sulovari et al.) is no longer considered human-specific, as homologous expanded alleles are now observed in both bonobo and chimpanzee haplotypes. (j) A newly identified HSE VNTR locus located within an intron of the *ART1* gene exhibits a bimodal copy number distribution and harbors distinct motif sequences compared to ancestral haplotypes in nonhuman primates.

Compared to previous reports^10^, we newly identified 2,756 human-specific TR expansions and corrected 1,113 loci that were previously misclassified, likely due to assembly limitations or technological biases. For example, a 4 bp STR in *NID2* (chr14:46237520-46237747) was incorrectly annotated as a human-specific expansion in earlier studies^10^, likely due to incomplete assemblies in nonhuman primates (Figure 5i). We also newly identified a 63 bp HSE VNTR in *ART1* (chr11:3699213-3720188), which exhibits a binomial copy number distribution in the human population (Figure 5j). VAMPIRE not only captured the variant motifs but also highlighted differences in both copy number and motif composition between orangutans, other nonhuman primates, and humans. These findings highlight both the improved sensitivity and biological relevance of VAMPIRE in identifying human-specific TR expansions.

Recent T2T NHP genomes revealed the presence of large subtelomeric heterochromatic caps (StSat, also known as pCht sequences), spanning approximately 1,193.7 Mbp, in *Pan* and gorilla genomes^3^. These regions are composed primarily of a 32 bp AT-rich satellite repeat, which is absent in human and orangutan genomes^24, 35^. The consensus motifs in *Pan* and *gorilla* differ by only a single nucleotide^36^ (A29->G), however, the extent and structure of motif variation within pCht sequences remain unclear. Using VAMPIRE, we systematically characterized motif variation in these regions and identified a total of 1,057 motifs across all pCht sequences in *Pan* and *Gorilla* (Supplementary Table 11). The 35 top-ranked motifs accounted for approximately 80% of the total pCht sequence length (Figure 6a, b). Notably, among the five top-ranked motifs, the first- and third-ranked motifs are distributed across both *Pan* and gorilla lineages (average 8.64%, 5.33% in gorilla and 12.06%, 8.02% in *Pan* respectively), whereas the second- and fifth-ranked motifs are largely restricted to gorilla (average 15.6%, 8.26% in gorilla and 0.68%, 0.98% in *Pan* respectively), and the fourth-ranked motif is predominantly found in *Pan* (2.66% in gorilla and 7.50% in *Pan*) (Figure 6d, e, Supplementary Table 12).

**Figure 6.**
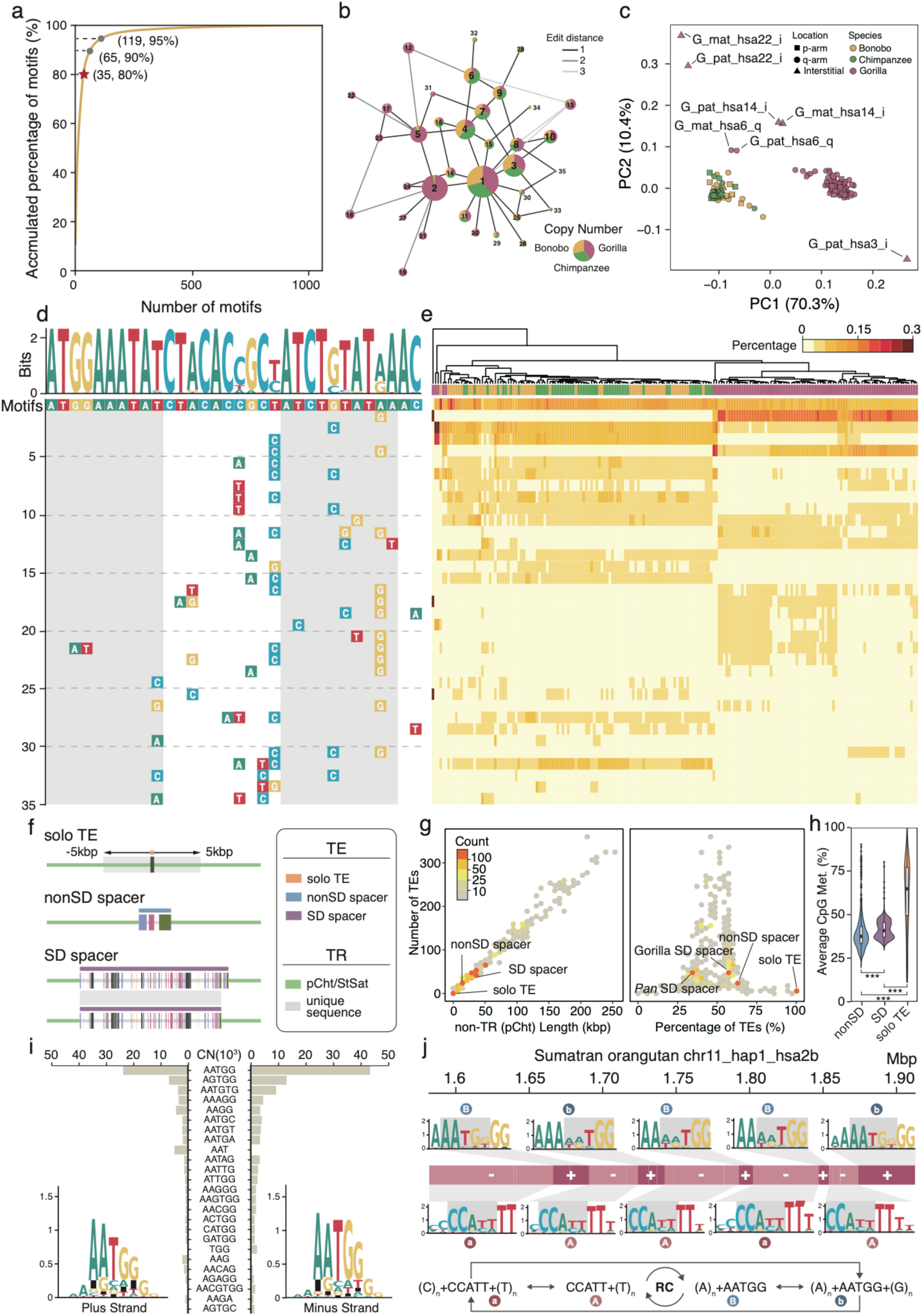
Discovering TEs, motif variation within subtelomeric heterochromatic caps of nonhuman African great apes. (a) Cumulative copy number percentage of 1,057 variant motifs identified by VAMPIRE. The top 35 motifs, collectively accounting for over 80% of total copies, were selected for downstream analysis. (b) Sequence similarity network of the 35 representative motifs. Each node represents a motif and edge color reflects the edit distance between motif pairs. (c) Principal component analysis of motif frequencies from 187 pCht/StSat sequences reveals two distinct clusters corresponding to *Pan* and gorilla, indicating different motif composition in two lineages. (d) Multiple sequence alignment and sequence logo of the motifs show that only four nucleotide positions are highly variable across the 32 bp consensus. (e) Heatmap of motif frequencies across sequences. (f) Schematic illustration of the three potential spacer types. As most TEs are located within 5 kb of one another, adjacent TEs within this distance were merged into single units, referred to as “non-SD spacers.” (g) Correlation distributions between spacer features: number of TEs vs. spacer length, and number of TEs vs. percentage of TEs. Identified SD spacers and non-SD spacers are labeled for reference. (h) Distribution of CpG methylation levels across the three spacer types. Both SD spacers and non-SD spacers exhibit CpG hypomethylation, suggesting similar epigenetic profiles. (i) Copy number distribution of the top-ranked HSATIII motif, showing that the plus strand of the AATGG motif is more conserved than the minus strand. (j) Example of an HSATIII region from orangutan chromosome 2b, demonstrating frequent inversions and diverse motif patterns of truncated TRs within these regions.

Principal component analysis (PCA) and hierarchical clustering revealed two distinct classes corresponding to *Pan* and *Gorilla* lineages, respectively. Interestingly, a subset of interstitial pCht sequences formed “outlier” groups (n=5), likely representing intermediate or ancestral forms between the two primary classes (Figure 6c). The previous studies focused on only two consensus motifs^36^, likely stemming from the limitations of TRF-based methods and the dominance of the A29→G site in distinguishing motifs (A29 vs. G29 frequencies: 46.09%/53.91% in gorilla and 94.52%/5.48% in *Pan*; (first-ranked and second-ranked)) (Supplementary Figure 16a, Supplementary Table 13). Our analysis reveals that these motifs are not strictly lineage-specific and are just dominant in each lineage.

In addition to confirming the previously reported SDs as spacers^3^, VAMPIRE further identified 191 solo and 677 clustered TEs (termed non-SD spacers) as additional spacers within the pCht regions, revealing a more complex structure than previously recognized (Supplementary Table 14). The non-SD spacers have an average length of approximately 22.6 kbp and a mean TE content of 50.5%, whereas solo TEs average only 434 bp in length, making them easily overlooked by TRF in the context of highly repetitive satellite-rich regions. (Figure 6f, g, Supplementary Figure 16b, c). Previous studies have reported lower

DNA methylation levels in SD spacers compared to adjacent pCht sequences. In this study, we observed that non-SD spacers exhibit significantly reduced methylation relative to SD spacers (Wilcoxon test, p<2.2e-16). While solo TEs also show lower methylation compared to pCht sequences (Wilcoxon test, p<2.2e-16), the variability across these regions is substantially higher. These findings suggest the existence of an additional class of spacer— non-SD spacers—embedded within pCht regions.” (Figure 6h, Supplementary Figure d-g). In all, the overall motif variation and organizational patterns within the pCht sequences are substantially more complex than previously appreciated.

Although pCht sequences are absent in orangutans, their subtelomeric regions differ from those of other primates and are collectively annotated as HSATIII^37^ (Supplementary Figure 17). Cytogenetic studies indicated that these regions do not display typical heterochromatic features^38^. To investigate this further, we analyzed the large satellite arrays on orangutan chromosomes 2a and 2b using VAMPIRE. Interestingly, we observed extensive TE insertions within the subtelomeric regions, disrupting the HSATIII satellite arrays. Moreover, these TR sequences are heterogeneous, consisting of multiple distinct motifs that frequently exhibit inversions and subclassifications relative to one another (Figure 6i, j). For both strands, there are two variation patterns (A and a for plus strand, B and b for minus strand). Pattern a and b have extra variants at the 3’ end of the consensus motif AATGG. These findings show that VAMPIRE is a powerful tool for uncovering the organization of complex, large-scale TRs, significantly advancing our understanding of satellite DNA architecture and evolution in non-model organisms.

## Discussion

TRs account for 4.9% to 13.0% of primate genomes^3^ (e.g., ∼3% in humans; 4.9% in Bornean orangutans and 13.0% in gorillas), with a significant portion localized in centromeric and subtelomeric regions—critical for chromosome segregation and genome structure maintenance^38, 39^. Increasing evidence has shown that TRs are not only associated with human diseases but also play a crucial role in genome size variation and contribute to interspecies genomic diversity^3, 40^. However, despite their importance, our understanding of their structure, variation, and molecular mechanisms in genome evolution, human diseases, and speciation remains limited.

To enable comprehensive analysis of TRs, we developed VAMPIRE, a tool that integrates motif annotation, variation detection, and structural motif decomposition. VAMPIRE automatically annotates motifs, while also identifying genetic variations in TR regions, such as non-TR sequences (e.g., TEs, SDs) and inversions. This integrated approach offers deeper insights into the composition and structural organization of TR motifs. In contrast, previous tools are often limited to generating consensus TR motifs and cannot detect SVs within large-scale TR arrays^15–18^.

Recent studies have highlighted the widespread occurrence of large-scale TR arrays across diverse organisms^3, 4, 41^, often correlating with adaptation. In contrast to earlier species-specific HMM approaches or database-dependent TR studies^20^, VAMPIRE provides a robust, versatile framework for investigating TRs in non-model organisms, significantly advancing our ability to explore their functional and evolutionary roles.

With the advancements of VAMPIRE, we focused on lineage-specific and expanded TRs in primate genomes. First, we uncovered previously overlooked diversity within the subtelomeric heterochromatin of nonhuman African great apes, highlighting motif turnover, variation, and diversity. Additionally, we deepened our understanding of orangutan subtelomeric repetitive regions by characterizing the relationships between motifs within these regions, rather than simply aggregating them as an HSatIII^42^. Further, we discovered recurrent inversions within the human centromeres^4^ and identified distinct inversion haplotypes in humans. The evolution of DNA satellite arrays is not limited to simple homogenization^43^; rather, diverse forms of genetic variation, including motif turnover, TE insertions, motif sequence inversions, and other rearrangements, frequently occur during TR evolution. These findings not only reinforce previous observations but also extend them, providing novel biological insights into the variation, structural organization, and evolutionary dynamics of TR motifs.

In summary, VAMPIRE streamlines TR analysis through a single-command execution, integrating multiple innovative algorithms to generate comprehensive TR profiles—from motif annotation and variation characterization to structural decomposition. As more complete genome assemblies become available, VAMPIRE will serve as a powerful tool for studying both simple and large-scale TRs across diverse organisms.

## Methods

### Algorithms implemented in VAMPIRE

#### Module 1: *De novo* motif finding

VAMPIRE segments the input sequence into overlapping windows (default: 5 kbp window size, 1 kbp step size) and processes them in parallel. Within each window, user-defined k-mers are generated and filtered based on a dynamic threshold: max{3, 0.01 × count of the most abundant k-mer}. A weighted *De Bruijn* graph is then constructed, where nodes represent k-mers and directed edges reflect observed k-mer transitions, with edge weights corresponding to transition frequencies. The graph is compacted by collapsing linear paths, and simple cycles (i.e., loops) are identified as candidate TR motifs. For each loop, the repeat count is estimated by the minimum edge weight along the cycle. Candidate motifs are ranked by estimated copy number, and the top motifs (default: 30) are retained as the initial motif set for downstream analysis.

#### Module 2: Motif iterative search

Both raw motifs and their reverse complements from the annotated motifs are inserted into a BK-tree to enable efficient approximate matching. Motifs are processed in descending order of abundance. For each motif, all possible rotations are queried against the BK-tree to identify a canonical form, ensuring consistent representation across redundant or equivalent sequences. By default, a match is defined as having ≥60% sequence similarity. If a match is found, the motif is replaced by its canonical counterpart. If no match is identified, the motif is treated as novel, assigned a canonical form based on its first appearance, and added to the BK-tree for subsequent queries.

#### Module 3: Motif alignment and chaining

This module generates the final annotation by aligning each candidate motif to the input sequence using edlib (v1.3.9). For every motif in the set, all potential matches are aligned, and matched regions are masked to prevent redundant hits. To determine the most likely arrangement of motifs across the sequence, we applied dynamic programming to identify the optimal annotation path. Each motif annotation is scored as: *length* ∗ *matchScore* − *dist* ∗ *distPenalty* − *lenDif* ∗ *LenDifPenalty* + *bonus*, where length is the motif length, dist is the edit distance between the motif and aligned sequence, and lenDif captures the difference in length, allowing the scoring system to prefer base substitutions over indels when the edit distance is equivalent. The recurrence relation is defined as:*dp*[*i*] = *max*(*dp*[*j*] + *score*[*t*], *dp*[*i* − 1] − *gap penalty*). Here, dp[i] represents the maximum cumulative score ending at position i, dp[j] represents the best score at the previous motif boundary, and score[t] is the score of motif t aligned from position j to i. A gap penalty is applied to penalize fragmented or sparsely distributed annotations. The final annotation corresponds to the highest-scoring chain, representing the optimal tandem repeat structure.

### Benchmarking in simulated data

We developed TR generator, a tool for simulating TR sequences with customizable parameters including repeat length, mutation rate, non-TR insertions, and motif composition. Using TR generator, we simulated 2,000 TR sequences with a 1% mutation rate, selecting consensus motifs from the VAMOS motif set^19^ based on the T2T-CHM13 assembly. To assess the impact of motif length, we divided the dataset into four subsets, each comprising 500 TR sequences with motif lengths of 1–6 bp, 7–19 bp, 20–49 bp, and ≥50 bp, respectively. For each motif, 1,000 copies were generated. To avoid redundancy, any simulated motif that could be represented in a shorter periodic unit (e.g., “TATTAT” as “TAT”×2) was discarded and replaced with a newly generated motif.

For TE insertion simulation, we randomly selected 1–3 TE sequences from the Dfam database (https://dfam.org/browse?clade=9443&clade_descendants=true) and inserted them at random positions within the sequence. A TE insertion was considered successfully detected if it met at least three of the following criteria: (1) The inserted TE motif was correctly identified; (2) The annotation achieved ≥80% sequence length accuracy; (3) The distance between the annotated breakpoint and the actual TE boundary did not exceed a defined threshold D, where D = *max*(15, 0.5 × *length of motif*) *bp*

For inverted sequence detection, we utilized the multiMotif function in TR generator to simulate tandem repeats containing both a motif and its reverse complement (e.g., “AATC” and “GATT”). An inversion event was considered successfully detected if it satisfied at least two of the following criteria: (1) Correct identification of both the original motif and its reverse complement; (2) Annotation accuracy ≥80% with respect to the full repeat sequence length.

For the mutation detection task, we constructed a motif set derived from the reference annotations and the outputs of the five evaluated tools:

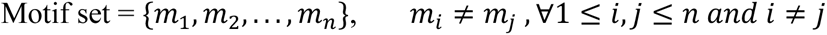

We then quantified the frequency of each motif in the results of each tool, representing them as n-dimensional vectors. To assess similarity between the predicted and reference motifs, we computed the Euclidean distance between these vectors, providing a quantitative measure of motif-level concordance across tools.

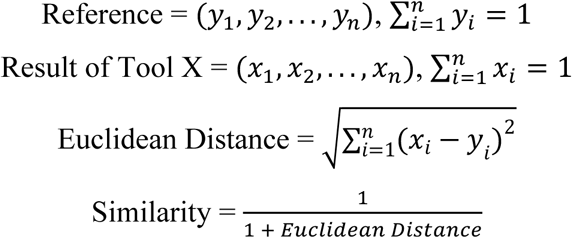

The parameters used for VAMPIRE, MotifScope (v1.0.0), TRF (v4.09.1), ULTRA (v1.0.3), and uTR (v1.0.0) are provided in Supplementary Table 1. Each benchmarking task was performed on three independently generated datasets, initialized with random seeds 665519, 184374, and 733241, respectively.

### TR comparison between VAMPIRE and TRF on T2T-CHM13 and HPRC

We developed a scan function to enable efficient TR detection across large genomic regions. This function partitions ultra-long sequences into discrete windows and executes VAMPIRE annotation in parallel using the default parameter set ‘-k 9’. For benchmarking, we also annotated the complete T2T-CHM13 assembly using TRF (v4.09.1) with the parameters “2 7 7 80 10 50 500 -h -l 20 -ngs”. To ensure high-confidence annotations, we excluded 1 bp motif loci and the loci with fewer than five total copies. A region was considered concordantly annotated by both tools if their respective annotations overlapped.

To map TR loci across HPRC haplotypes, we extracted 1 kbp flanking regions on both sides of each TR locus as anchor pairs and used LiftOver (ucsc) to project their coordinates across 94 assemblies from the HPRC^44^. To ensure accurate annotation of haplotypes with low copy numbers, we implemented a two-step strategy: first, we constructed a motif set using the command ‘-k 11’; second, we re-annotated the sequences with this motif set using parameter ‘--no-donovo –forcè. To evaluate annotation accuracy, we employed the evaluate function in VAMPIRE to calculate the mean edit distance between annotated motifs and the corresponding reference sequences.

### Centromere annotation, similarity and expansion Analysis

To benchmark the performance of VAMPIRE and ModDotPlot^31^, we annotated all 24 (22 autosomes and XY) active HOR regions in the T2T-CHM13 assembly. Monomer-level annotation was performed with command: vampire anno -k 13, and sequence similarity was computed with VAMPIRE using a non-overlapping window of 60 motifs. Sequence similarity (percentID_by_events) was defined as the sum of matched bases (M), mismatches (X), and insertion (I) and deletion (D) events: similarity = 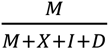. To focus specifically on single-nucleotide divergence, we set MIN_LENGTH and MAX_LENGTH to zero, effectively excluding indels and simplifying the similarity calculation to similarity = 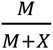.

To control for potential bias introduced by varying window sizes, ModDotPlot^31^ was run using a fixed 10 kb window. Concordance between the two tools was evaluated by computing the Pearson correlation coefficient across all window pairs with ≥90% reciprocal overlap. To assess structural similarity, similarity heatmaps from both methods were normalized using Min-Max scaling, aligned to identical dimensions, and compared using the Structural Similarity Index Measure (SSIM), implemented via scikit-image.

For the analysis of the human chromosome 1 centromere (CEN1), we selected 31 samples from the HGSVC3 dataset ^32^ that contained fully assembled CEN1 regions. Similarity heatmaps were visualized using a customized version of the StainedGlass^33^ plotting scripts, with additional tracks to display TR and TE annotations, as well as strand orientation. TR/TE annotations were generated using RepeatMasker^42^ (v4.1.4) with the parameter ‘-species primates’, and strand information was extracted directly from the VAMPIRE annotation files.

### Human-specific expanded (HSE) TR and burst HSE identification

We used T2T diploid genome assemblies from bonobo, chimpanzee, gorilla, Sumatran orangutan, and Bornean orangutan^3^, and a T2T haploid genome from the crab-eating macaque^45^ as outgroup species. To capture representative human population diversity, we selected 10 individuals from the HPRC dataset^44^, including 5 African (HG02257, HG02717, HG03453, NA19240, NA20129), 2 American (HG01123, HG01952), 2 East Asian (HG02080, HG00673), and 1 South Asian (HG03492) samples. We used the 1 kbp flanking sequences surrounding each tandem repeat locus as anchors and applied liftOver to map these regions across assemblies. After that, we extracted the corresponding sequences and merged them into a fasta file for each locus. To fully capture motif variation across species, we used consensus motifs to annotate and extract all mapped sequences from the initial annotation results, generating a unified motif set across species. Consequently, we annotated each sequence using this combined motif set to ensure consistent motif annotation across samples. Total copy number and copy number for each motif are calculated for each locus in each genome. Loci supported by fewer than 10 human assemblies and 5 nonhuman primates assemblies are excluded from downstream analysis. We then applied the filtering procedure described in Sulovari et al.^10^.

To identify human-specific expanded tandem repeats (HSEs), we performed a nonparametric one-tailed Wilcoxon rank-sum test to assess differences in TR lengths between human and NHP groups. Multiple testing correction was applied using the False Discovery Rate (FDR) method, retaining loci with q-values ≤ 0.05. To further quantify divergence in TR variation, we calculated the VST statistic for each locus using the following formula:

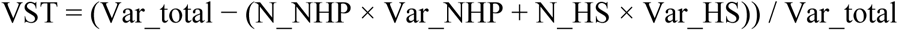

where: Var_total is the total variance across all samples, Var_NHP and Var_HS are the variances within the human and NHP cohorts, respectively, N_NHP and N_HS denote the number of haplotypes in each group.

To establish a threshold for significant divergence, we applied the extreme distance estimator combined with the Chebyshev confidence interval to identify the first inflection point in the VST probability distribution. This threshold was determined to be at VST = 0.50, with loci showing VST ≥ 0.50 are considered strong candidates for human-specific repeat expansions. Finally, we required that the minimal difference in TR copy number—between the shortest human and longest NHP alleles—exceed 5 repeat copies.

To quantify the copy number dynamics of human-specific expansion (HSE) locus, we defined an HSE index based on copy number (CN) statistics. The index was calculated as:

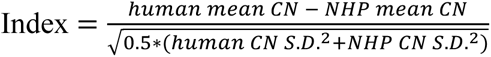

Here, S.D. is the abbreviation of standard deviation. To identify the most prominent inflection point in the distribution of HSE indices, we computed the kernel density estimate (KDE) of the HSE index values and derived the cumulative density function (CDF). The slope of the CDF was calculated to locate the point of maximum change. The x-coordinate corresponding to the maximum slope was used as the empirical threshold to distinguish loci with pronounced HSE signal. We defined HSE locus with index greater than 4.63 as burst locus, indicating a rapid and substantial expansion in humans, rather than a gradual or merely accelerated process.

### pCht/StSat and HSATIII variants identification and composition analysis

Recent studies have identified the pCht sequences—characterized by a 32 bp TR motif expansion interspersed with SD insertions serving as spacers—in the genomes of chimpanzees, bonobos, and gorillas. To systematically characterize these pCht regions, we first conducted a whole-genome scan across three T2T NHP assemblies. TR sequences exhibiting more than four base differences relative to the consensus pCht motif (ATGGAAATATCTACACCGCTATCTGTATAAAC), corresponding to a minimum similarity threshold of 87.5%, were excluded. To account for the presence of inserted SD spacers, adjacent TR regions separated by less than 1 Mbp were merged into single contiguous loci for downstream analyses.

Each region was named following the format species-haplotype-chromosome-location, reflecting the species, haplotype, corresponding human chromosome, and genomic coordinates. Regions were further classified into three categories: p-arm, q-arm, and interstitial. We then employed VAMPIRE for motif annotation using the parameters: vampire anno -k 13 -n 50. To streamline downstream analyses and improve interpretability, we calculated the cumulative repeat counts for each motif and selected the 35 most abundant motifs, which together accounted for 80% of the total pCht sequence counts. These representative motifs were subsequently used to annotate pCht sequences with the parameters ‘vampire anno --force --no-denovo -m’.

For motif sequence comparison, pairwise alignments were performed among the 35 selected motifs, and their relationships were visualized as a graph where each node represents a motif sequence and each edge corresponds to an alignment. Only the highest-scoring match for each node was retained in the final graph. To characterize base composition, sequence logos were generated based on VAMPIRE annotation results using the logo function of VAMPIRE. Principal component analysis (PCA) was conducted on motif frequency profiles across sequences using the scikit-learn library.

We linked transposons annotated by RepeatMasker^42^ (v4.1.4) within 5 kbp and classified them into three types based on the number of TEs and sequences—solo TE, nonSD spacer and SD spacer. Solo TE is the only transposon without any other TEs in 5 kbp flanking regions. SD spacer is clustered TEs whose sequences are part of segmental duplications. And nonSD spacer is clustered TEs but not belongs to any segmental duplications.

We used the 5 bp consensus motif AATGG to decompose the HSATIII sequences with the default parameters. Adjacent regions on the same strand were merged into continuous segments, which were then analyzed for motif variation using sequence logo plots.

## Acknowledgments

We thank the HPRC, APC, and Primate T2T Consortium for providing the long-read primate genome assemblies. The computations in this study were run on the Siyuan-1 supported by the Center for High Performance Computing at Shanghai Jiao Tong University.

## Authors’ contributions

Y.M. conceived the project. Z.Y. and S.Z. developed and validated the tool. Z.Y. performed all data analysis. Z.Y., G.A.L., and Y.M. performed centromere analysis. Z.Y. and Y.M. drafted the manuscript. All authors read and approved the manuscript.

## Data Availability

The code for VAMPIRE is available on GitHub (https://github.com/Zikun-Yang/VAMPIRE) with an MIT license. The assemblies or genomic sequences used in this study (except HGSVC3 and HPRC dataset) can be found in GenBank with the accession number GCA_009914755.4, GCA_028858775.2, GCA_028885625.2, GCA_028885655.2, GCA_029281585.2 and GCA_029289425.2.

